# Tracking Action in Cross-Reality Mark Test Reveals Developing Body Representation Among Toddlers

**DOI:** 10.1101/2021.10.08.462966

**Authors:** Michiko Miyazaki, Tomohisa Asai, Norihiro Ban, Ryoko Mugitani

## Abstract

Research on multisensory integration—particularly visuo-proprioceptive integration—has renewed interest in how body representation develops in early childhood. In the past decade, researchers have attempted to adapt body-illusion tasks such as the rubber hand illusion for young children, yet difficulties with verbal reports and inconsistent behavioral responses limit their usefulness for toddlers. At the same time, children naturally begin to recognize their mirror image as self during the early toddler period, suggesting that mirror-based localization may offer a developmentally appropriate window onto emerging body representation. However, the classic mark test is not suitable for repeated measurements. We developed Bodytoypo, a cross-reality body-part localization task that projects a life-size self-image onto a screen and overlays AR markers on multiple body locations using real-time skeleton detection. Toddlers touched the corresponding location on their actual body, allowing quantitative evaluation of localization accuracy. Thirty 2- and 3-year-olds participated. We recorded first-touch outcomes, touch error, reaction times, and hand trajectories. Conventional coding was combined with an automated statistical analysis to estimate factors predicting touch error. Results showed that Bodytoypo is feasible for repeated use with toddlers and provides reliable indices of body-part localization. Automated statistical analyses identified several reaching strategies that appeared to reflect differences in localization precision. These findings suggest age-related changes in how visual information and motor prediction are weighted during visuo-proprioceptive integration. Bodytoypo offers a promising behavioral index for clarifying how sensorimotor information (body schema) and spatial understanding of body parts (body topology) jointly contribute to the early development of body representation.

## I. INTRODUCTION

RESEARCH on body representation has expanded alongside developments in embodied cognition, with concepts such as *body schema* and *body image* providing influential frameworks for explaining diverse cognitive and motor phenomena [1]–[5]. However, the traditional dichotomy that treats the body schema as an implicit, sensorimotor representation and the body image as an explicit, visually driven representation has proven insufficient to account for the variety of impairment patterns reported in clinical and behavioral studies [6]–[8]. Schwoebel and Coslett [8] showed—through principal component analysis of task performance in patients with unilateral brain damage—that body image comprises at least two distinct components: *body topology*, representing spatial relations among body parts, and *body semantics*, referring to knowledge of body-part names and functions. Together with the body schema, these findings support a tripartite structure of body representation, yet how these components emerge and differentiate during early development remains largely unknown.

Work with infants and toddlers has accumulated findings on body semantics and body topology, including discrimination of body-part location violations [9], acquisition of body-part labels [10]–[12], and pointing accuracy to different body locations [13], [14]. Nevertheless, these findings primarily address visually guided and vocabulary-based aspects of development, offering limited insight into when the sensorimotor body schema emerges and how it becomes coordinated with visuospatial based body topology. Consequently, the developmental correspondence of the tripartite framework—body schema, body topology, and body semantics—remains unclear, posing a central challenge for constructing a developmentally grounded account of body representation.

Recent work examining infants’ motor responses offers clues for addressing this gap [15]–[17]. Somogyi et al. [17] tested 43 infants aged 3–6 months by applying vibrotactile stimulation to multiple body locations and closely observing the resulting movements. Only five infants were able to remove the vibrator themselves, but qualitative analyses of the remaining infants revealed location-specific motor responses that became more differentiated with age. These findings indicate that rudimentary body-topological distinctions can be captured through nonverbal behavioral measures and suggest that interactions between the sensorimotor body schema and visually based body topology may already be observable in early infancy. However, the limited number of successful trials and the reliance on qualitative analyses underscore the need for methods that can directly and quantitatively evaluate how *visuo-proprioceptive* cues are integrated during early development.

Among experimental paradigms used to investigate body representation, visuo-tactile and visuo-proprioceptive illusion methods—most notably the rubber hand illusion (RHI)—have played a central role [18]. In the RHI, synchronous stroking of a visible fake hand and the participant’s hidden real hand induces a sense of ownership over the fake hand, enabling researchers to examine how visual, tactile, and proprioceptive cues are integrated. However, because the RHI typically relies on verbal reports to assess the induced illusion, it is difficult to apply to toddlers, who often show affirmative-response biases. In addition, proprioceptive drift—the primary non-verbal behavioral measure used in RHI studies—has recently been shown to be highly inconsistent across studies according to meta-analytic evidence, further limiting the suitability of the RHI for early developmental populations [19]–[22]. In contrast, the task in which individuals locate a mark on their own body by relying on information from their mirror image—the mark test—was originally proposed as an index of mirror self-recognition [23], [24]. However, aligning one’s visual mirror image with one’s physical body requires the same type of visuo-proprioceptive correspondence operations that underlie body part localization. From this perspective, infants’ behavior in mirror-based situations can offer a valuable window into the early development of body representations. Indeed, prior work has shown that the likelihood of localization errors varies substantially depending on where the secretly applied mark is placed (e.g., forehead vs. nose) [25], [26], suggesting that error patterns in the mark test reflect developmental differences in body representation. Nevertheless, because the mark test requires applying a physical mark to the child’s body, once children become aware of the procedure, repeated measurements become difficult, thereby limiting its usefulness in trial-by-trial analyses [27]. Consequently, both the RHI and the mark test face critical constraints for capturing the dynamics of early visuo-proprioceptive integration in a repetitive and quantitatively robust manner.

The present study aims to develop an augmented-reality (AR)–based body-part localization task for toddlers and to evaluate visuo-proprioceptive integration through the analysis of reaching trajectories. By projecting virtual markers onto children’s bodies, the task elicits natural touching and reaching behaviors, enabling nonverbal and repeatedly measurable assessments beyond the constraints of prior methods. Analyzing visuo-motor trajectory patterns, together with localization errors and reaction times, allows us to describe when and how the body schema and body topology begin to integrate and how this integration varies across body regions. This study introduces a novel behavioral index of multisensory body representation and provides foundational evidence for developmentally situating the tripartite framework.

## II. Cross-reality MARK TASK: Bodytoypo

### A. Apparatus

We developed a body part localization task using the image processing library Openpose [28] and augmented reality (AR) (see Fig. 1 and Fig. S1A). An image of each participant’s full body was recorded using a USB camera (Logicool C920) and projected life-size onto a screen (KIMOTO RUM60N1) via a projector (Epson EB 485WT). Virtual marks were sequentially displayed on 30 parts of the participants’ bodies. The 2D coordinates of each body part were detected in real-time using Openpose on a GPU machine (Mouse computer NEXTGEAR i690PA2), and AR characters (favored cartoon characters of toddlers) were displayed as marks on the coordinates of the target body parts. The net delay for transmission and image processing was approximately 10 frames (0.33 seconds), and the AR overlay processing speed ranged from 15 to 22 fps (best-effort format). Participants stood on a designated spot 115 cm from the screen, and the camera angle was adjusted so their full body was reflected on the screen. Each participant’s behavior was recorded using a capture device (Avermedia AVT-C878) on a stand-alone PC (Galleria QSF1070HGS). The recorded videos were later used for offline manual coding and movement analysis. We named this system “Bodytoypo.” Bodytoypo was an interactive cross-reality mark test that crosses the boundary between real and virtual, allowing participants to *touch* AR marks.

**Fig. 1.**
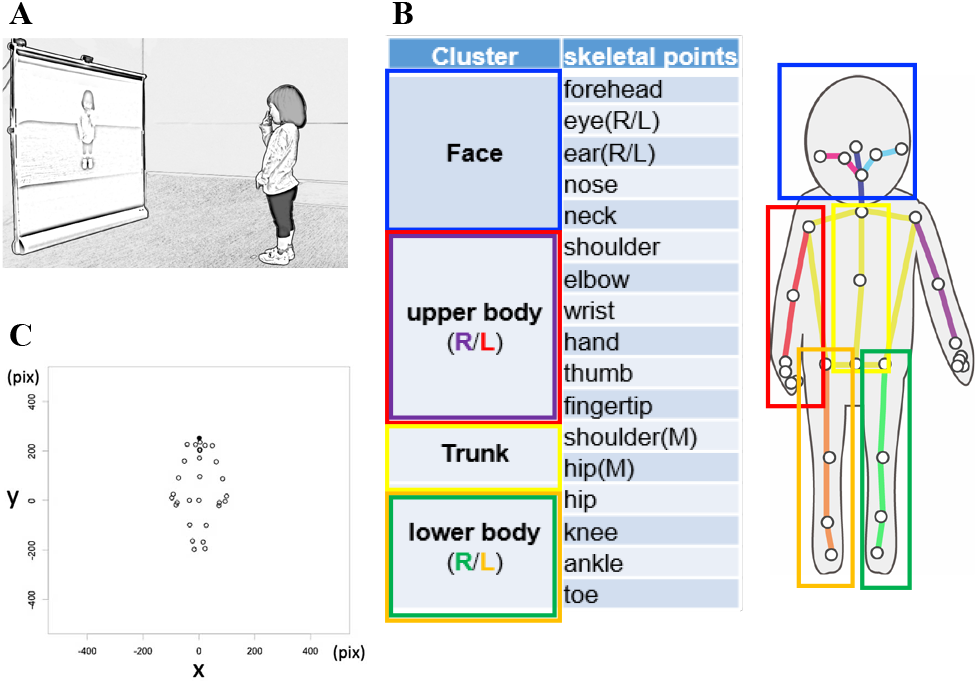
Real-time estimation of body parts for the cross-reality mark test (“Bodytoypo”). Note: (A) Online presentation for participants touching the AR mark in the current study. (B) The original input image (frame of the video) with the estimated bone (30 parts) and six body clusters using automated statistical analysis. R/L/M in the table indicate right, left, and midline, respectively. (C) The depicted output for x-y coordinates of each body part (30 fps) for the following offline motion analysis. The location of the center of the hip (Chip, 22) was always fixed at the origin (0,0) as the reference point.

### B. Body Part Localization Task (Bodytoypo)

The experimenter manually displayed the AR mark on the participant’s life-size self-image and observed whether the participant could correctly touch (or point to) the actual body part corresponding to the mark’s location. The AR marks were displayed individually on 30 body parts (upper forehead, forehead, eye (right/left, hereafter R/L), ear (R/L), nose, cheek (R/L), mouth, chin, collarbone (R/L), shoulder (R/L), upper arm (R/L), elbow (R/L), navel, forearm (R/L), hand (R/L), thigh (R/L), and big toe (R/L)). Note that the 30 body parts shown in Bodytoypo were based on the Openpose definition, which slightly differed from those in the independent offline analysis (see Fig. 1B, Fig. S1B, C). The marks were arranged in a random sequence to avoid adjacent body parts. These sequences were presented in forward and reverse directions. The presentation order was counterbalanced among participants. If the participants could touch the correct body part with the AR mark, the mark disappeared with a fun audio-visual reward presented by the experimenter’s manual keyboard control.

### C. Analytical Strategy

The present study addressed two main research questions: (1) whether Bodytoypo can provide reliable behavioral indices of body-part localization and reaching trajectories, and (2) whether these indices can capture developmental changes in visuo-proprioceptive integration during early childhood.

To answer these questions, we focused on the period from the appearance of an AR mark to the toddler’s first touch. The analysis combined conventional manual coding and statistics-based automated movement analysis. Manual coding yielded trial counts, accuracy for each body part, and reaction times, allowing us to characterize which body parts were easier or harder to localize. Although manual coding can be expressed either as accuracy or error rates, we adopted an error-based approach consistently across all analyses. This decision reflects the fact that the automated statistical analysis relies on a continuous distance measure—the pixel distance between the target and the first touch—to quantify performance because binary accuracy cannot capture the magnitude of mislocalization. Therefore, all evaluations in this study focused on errors rather than accuracy. Automated analysis was then used to quantify movement dynamics. First, each trial was divided into a *latency* period and a *reaching* period based on temporal changes in the spatial relationship between the acting hand and the target (see *Analysis Pipeline*). Second, to characterize the sensorimotor coordinates involved in touching the AR mark, we calculated whole-body associations during these two periods. Finally, the correlation matrix was vectorized and entered into the regression analysis with reaction time and age, to evaluate how well these variables predicted toddlers’ touch errors. These strategies allowed us to evaluate whether movement trajectories provide meaningful signatures of multisensory body representation.

## III. Methods

### A. Participants

A total of 36 toddlers living in Japan, aged 2.5 to 3.5 years, participated in the study. The sample size was determined based on previous studies on body representations [10], [17]. The study was conducted in accordance with the recommendations of the Otsuma Women’s University Life Sciences Research Ethics Committee, and written informed consent was obtained from all parents. The protocol was also approved by the same ethics committee (2019-012). The final sample consisted of 30 toddlers (mean age = 34.7 months, range: 28–44 months, SD = 4.49), including 18 2.5-year-olds (11 females) and 12 3.5-year-olds (9 females). Six participants were excluded (attrition rate = 16.7%) due to fussiness and shyness (n = 5) or technical issues with the equipment (n = 1).

The final dataset for the automated statistical analysis included 607 trials (2.5-year-olds: n = 310; 3.5-year-olds: n = 297) out of a total of 852 manually coded trials. Because our objective was to evaluate localization errors and the motion trajectories associated with the toddlers’ first touch, 245 trials (28.7%) were excluded as unsuitable for trajectory analysis. The primary exclusion criterion was an excessively long reaction time (greater than 3 seconds, approximately 3 SD above the mean). Such trials were largely affected by behaviors unrelated to task-relevant motor responses—for example, the child not noticing the target mark, showing reduced motivation to touch, becoming distracted during play, or temporarily disengaging from the task.

### B. Tasks and Procedures

#### Bodytoypo Task

The experimenter first explained the study to the parents to obtained informed consent. The experimenter and the toddlers then developed rapport through free play. The experimenter demonstrated ten Bodytoypo trials. During the demonstration, she intentionally touched an incorrect location and informed the toddlers that incorrect touches would not yield a reward. When the participants were standing straight and still in front of the screen, an AR mark appeared with a beep sound. The experimenter encouraged participation by rhythmically presenting the AR marks. If the participants did not respond, the experimenter re-presented the mark or skipped the trial. To maintain motivation, additional trials involving touching the easily accessible nose were inserted as needed. Aside from these additional nose trials, a maximum of 30 trials (30 body parts) were administered to each toddler. The task lasted approximately five minutes.

#### Questionnaire of Body Part Vocabulary

After the Bodytoypo task, parents completed a questionnaire assessing their child’s production and comprehension of 60 body parts, including human-specific and animal-related parts (e.g., tail, wings, horns) in Japanese.

## IV. Results

### A. Validity of the Body Part Localization Task

#### Maintenance of Participants’ Motivation and Overall Error Rates

The mean numbers of executed trials were 27.50 (SD = 5.371) and 29.75 (SD = .452) for the 2- and 3-year-olds, respectively. Most participants completed 30 trials regardless of age (*t*(28) = -1.439, *p* = .161, *d* = -.522). This result was consistent with previous findings [26] and suggests that participants’ motivation was successfully maintained throughout the task. Furthermore, the pattern of results was also consistent with a positive effect of gamification, including animated feedback and sound effects.

The overall first-touch error rates were 39.9% (SD = .132) for 2-year-olds and 35.0% (SD = .095) for 3-year-olds. The age groups did not differ significantly (*t*(28) = 1.098, *p* = .282, *d* = .398). Figure 2 summarizes these error rates for each age and body part (Fig. 2A) and for each mark location (midline, left side, right side; Fig. 2B). To examine the role of mark location, we grouped body parts according to whether the mark appeared on the midline, left side and right side. The main effect of age was not significant (*F*(1,28) = .830, *p* = .370), nor was the interaction between age and mark location (*F*(2,56) = .954, *p* = .391). However, the main effect of mark location was significant (*F*(1,28) = 35.647, *p* < .01, partial *η*^*2*^ = .560). Holm-adjusted comparisons indicated that midline error rates were significantly lower than those on the left or right (midline–left: *p* < .01, *d* = –1.635; midline–right: *p* < .01, *d* = –1.338), whereas left–right differences were not significant (*d* = 0.385).

**Fig. 2.**
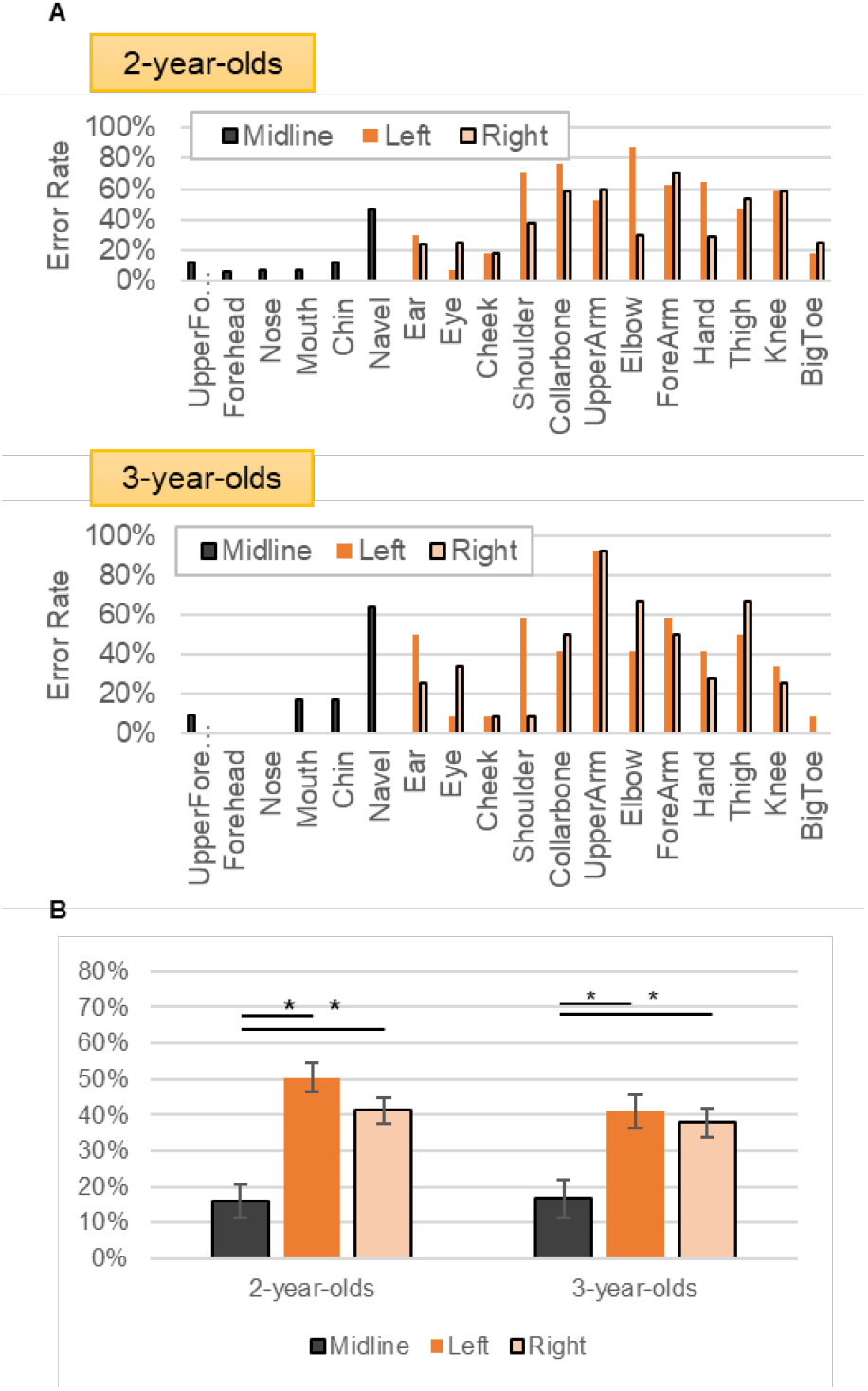
First-touch error rates in the Bodytoypo task. Note: (A) Mean error rates for all 30 body parts by age group. Body parts located on the face showed consistently low error rates for both ages. (B) Mean error rates by mark-location category (midline, left, right) for each age group. Error rates for midline body parts were significantly lower than those for parts on the left or right sides.

#### Localization Errors and Vocabulary Acquisition

We further analyzed localization error rates by categorizing the 30 body parts according to four bodily functions: joints, movable parts, face parts, and lexically specified body parts (Fig. 3). A two-way ANOVA (age × bodily function) revealed significant main effects for all categories (joints: *F*(1,28) = 12.259, *p* = .002, *MSe* = .021, partial *η*^*2*^ = .305; movable parts: *F*(1,28) = 25.507, *p* < .01, *MSe* = .013, partial *η*^*2*^ = .149; face parts: *F*(1,28) = 113.637, *p* < .01, *MSe* = .015, partial *η*^*2*^ = .802; lexically specified parts: *F*(1,28) = 58.481, *p* < .01, *MSe* = .017, partial *η*^*2*^ = .676). The interaction between age and bodily function was significant for joints (*F*(1,28) = 4.898, *p* = .035, partial *η*^*2*^ = .149) and marginally significant for movable parts (*F*(1,28) = 4.026, *p* = .055, partial *η*^*2*^ = .126). These results suggest that the accuracy of first-touch localization improves between ages 2 and 3 for joints and movable parts.

**Fig. 3.**
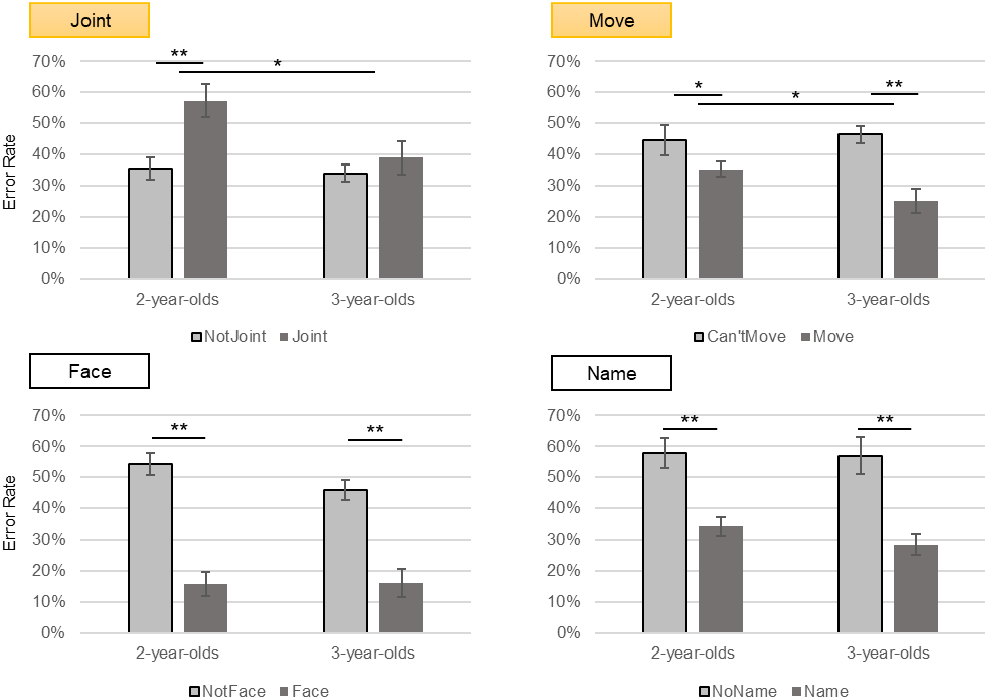
First-touch error rates categorized by bodily function. Note: The 30 body parts used in the Bodytoypo task were classified according to four functional categories: joint vs. non-joint, movable vs. non-movable, face vs. non-face, and lexically specified vs. non-specified. Significant main effects were observed for all four categories. Interactions with age emerged for the joint and movable categories, indicating age-related improvements in localizing joints and movable parts.

Parental reports of body-part vocabulary revealed that children produced and comprehended an average of 28.7 words (SD = 10.599) at age 2 and 40.8 words (SD = 14.640) at age 3 out of the 60 items queried (Fig. S2C). This age difference was significant (*t*(28) = –2.632, *p* = .014, *d* = –.955). Partial correlation analyses controlling for age in months indicated no significant association between total body-part vocabulary and Bodytoypo error rates (Fig. S2A). Likewise, no significant relationship was found between vocabulary for the body parts used in Bodytoypo and the corresponding error rates (Fig. S2B).

### B. Exploratory Analysis of Movement-Based Evaluation Metrics

#### Analysis Pipeline

The overall pipeline in the statistical analysis is shown below. The necessary process was first; the change of point detection as each trial (from the mark appearing to the first touch) comprised two periods: response latency and reaching movement (Fig. 4). Therefore, the change point was estimated using the machine-learning R library “changepoint” (https://github.com/rkillick/changepoint/) over the time series of variance among distances between the used hand and other body parts (Fig. S4). During a reach, these distances change drastically compared with those during latency. This automated estimation (separation of latency and reaching) was further validated during a process conducted later (Fig. 5ABC and 6AE). Once we distinguished them, the whole-body associations were calculated as a temporal-correlation matrix during those two periods (Fig. 5), indicating sensorimotor coordinates for touching the AR mark. This summarized matrix exhibits a relative similarity of trajectory among body parts so that the original configuration (physical body) was partly recovered from the matrix through multidimensional scaling (MDS) projection during latency (Fig. 6A) rather than during reaching (Fig. 6E). Furthermore, this correlation matrix was vectorized and entered into the regression analysis (as inputs) to predict toddlers’s touch errors (as outputs) (Fig. 6BD). During each period, body part associations were visualized at a glance in terms of the x-y joint dendrogram (tanglegram, Fig. 6CF). Lastly, all the variables were put into a linear mixed model (all-in-one statistical model) as continuous variables, to predict touch error (Fig. 7A). In summary, we established that 2- to 3-year-olds’ sensorimotor representation can be depicted where the feedback or feedforward weight for touching seems to be adaptively changing.

**Fig. 4.**
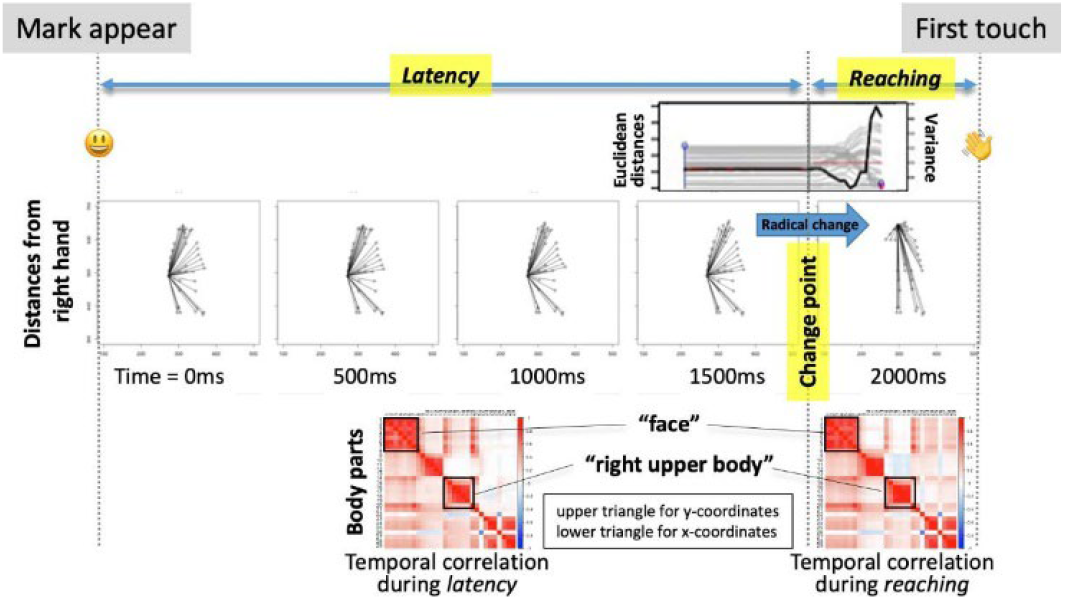
Schematic illustration for each trial. Note: Each trial (from the mark appearing to the first touch) has two periods: response latency and reaching duration. This was defined by the detected change point over the time series of variance among distances between the used hand (in this example, right hand) and other body parts (the upper panel or supplementary Figure S4). During reaching, these distances dramatically change compared with those during latency (middle successive panels). The association among body parts (temporal correlation of x- or y-components) was calculated for each period (lower two panels), where some “clusters” are observed on the diagonal (e.g., “face” cluster). This correlation matrix was further vectorized and used in the statistical analysis as inputs for predicting participants’ touch errors as outputs.

**Fig. 5.**
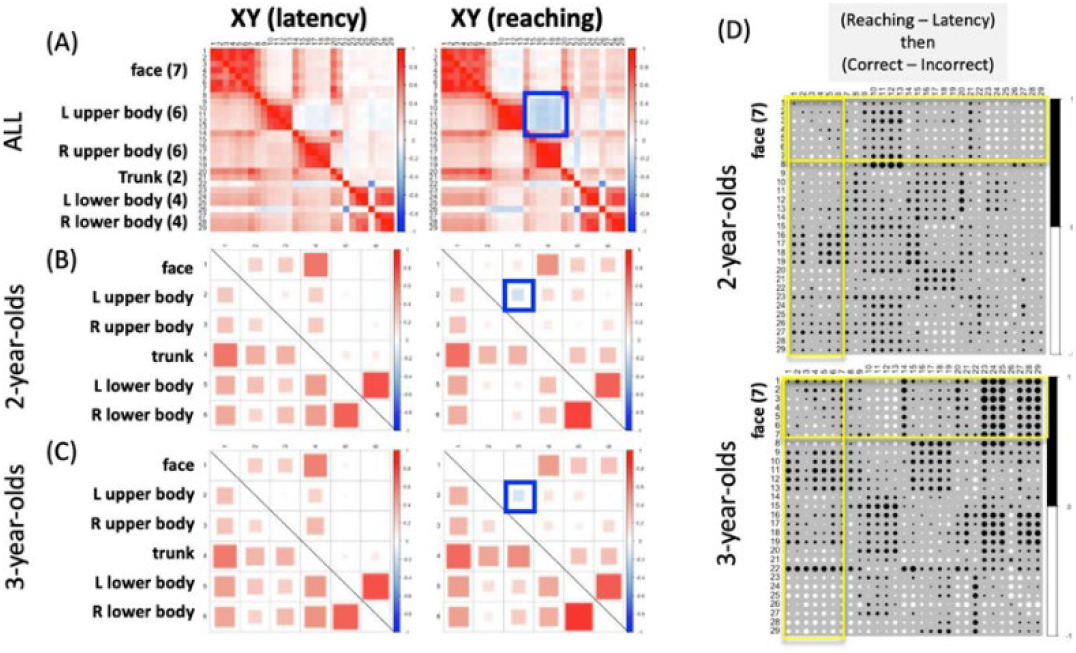
Temporal Correlation Among Body Parts for 2- or 3-year-olds. *Note*. Temporal correlation matrices for body parts during latency or reaching. The upper triangles are for x-, whereas the lower triangles are for the y-coordinates. Body parts (A) or clusters (B, C) were positively correlated in general, except for L-R upper-body relations during the reaching period (blue squares). This negative correlation in the x-coordinate suggests bimanual coordination in reaching behaviors that could validate the change point estimation. Both age groups behaved similarly in terms of the association among body clusters (B, C) as well as of the reaching trajectory (see Fig. S5). (D) However, the face behaved oppositely between 2- and 3-year-olds if we look at the subtracted matrix between correct and incorrect trials (see the main text for the detailed discussion).

**Fig. 6.**
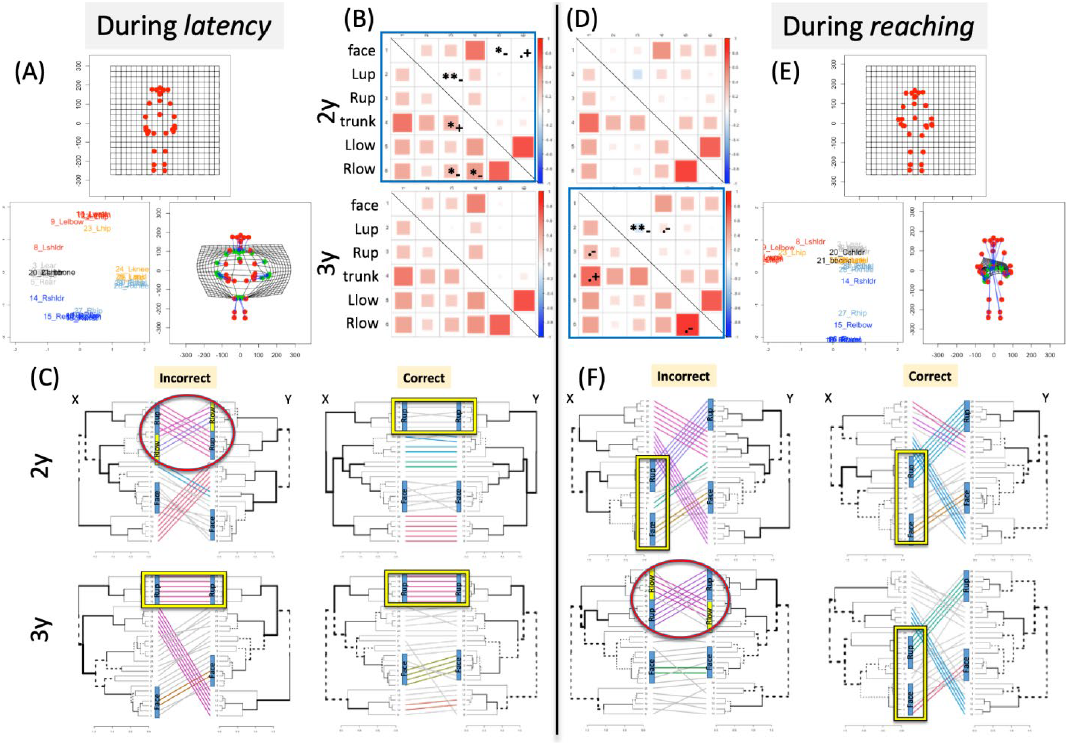
Body Parts Associations for Configuration (AE), Prediction (BD), and Coordination (CF) *Note*. The temporal correlation matrices for latency (A) and reaching (E) were analyzed to recover the physical body parts configuration through MDS projection and the Procrustes transformation. However, the reaching matrix did not globally recover the configuration due to the inclusion of various movements. The latency matrix for 2-year-olds can statistically predict the later touch errors (B) while the reaching matrix for 3-year-olds can predict the later touch errors (D). During the latency period, the incorrect tanglegram for 2-year-olds was characterized by a right upper and lower parts association (C). During the reaching period, the same association was observed for the incorrect tanglegram for 3-year-olds (F). See the main text for more discussion.

**Fig. 7.**
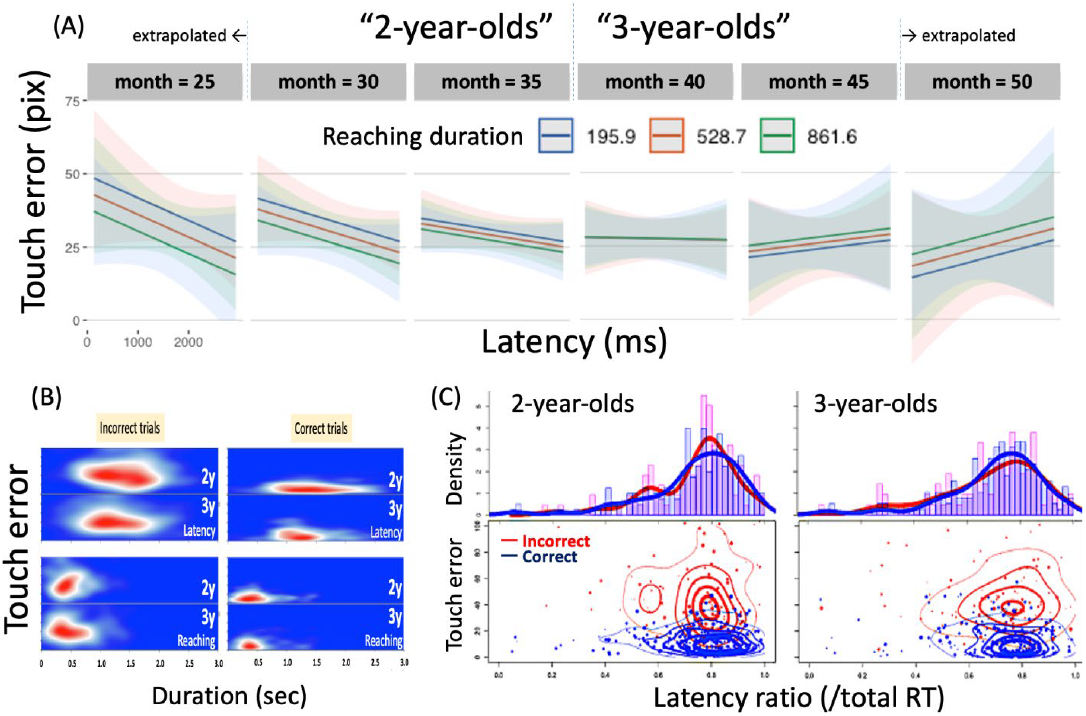
Estimated Parameters for Predicting the Touch Error Regarding Age in Months. *Note*. (A) A linear mixed model fitted using the Akaike Information Criterion (AIC), to determine the most informative inputs. The model examined how latency, reaching duration, and age (in months) could predict touch errors. Interestingly, the slopes of these predictors gradually reversed as children got older. (B) Heatmaps to visualize the distribution of latency/reaching duration and touch errors across different ages and correct/incorrect trials. (C) The contour plots of (B) for comparison (lower) and probability distributions for latency ratio. In summary, two- and three-year-olds have opposite touching strategies in terms of feedback/forward weights.

Furthermore, the effectiveness of the automated analysis was evaluated by applying the findings from the statistics-based automated analysis to the manual analysis, from a qualitative perspective (Fig. S3).

#### Body-Parts Association During Latency and Reaching

Figure 4 summarizes the procedure of each trial and Figure 5 suggests the averaged temporal correlation among 30 body parts (29 parts except for the reference). The upper triangles represent x-, while the lower triangles represent y-coordinates where some “clusters” are observed on the diagonal (e.g., “face” cluster). The body parts (Fig. 5A) or clusters (parts-averaged, Fig. 5BC) were generally positively correlated, except for L-R upper body associations during the reaching period (blue squares). This negative correlation in the x-coordinate suggests bi-manual coordination in reaching behaviors, which could validate the change point estimation (see Fig. S4). Both age groups behaved similarly in terms of these correlation matrices (Fig. 5BC) as well as of the reaching trajectory in 2D (Fig. S5). However, the “face” (or head) behaved oppositely between 2- and 3-year-olds in the case of the subtracted matrix between correct and incorrect trials (yellow parts in Fig. 5D for reverse monochrome patterns) or the cross-correlation between the face and the hand (Fig. S7A). We observed what happened to those participants as follows.

#### Correlation Matrix for Configuration, Prediction, and Coordination

Figure 6A shows the recovered body configuration from the physical positions (averaged x-y coordinates of all trials, the upper panel) to the MDS projection (based on the averaged temporal correlation matrix among body parts and the lower-left panel) during latency. The comparison between them (through the Procrustes transformation in the lower right panel) suggested that the association between body parts during latency is relatively stable so that we can see the original body configuration. However, it is horizontally biased as the hand moved horizontally with ease, even during latency. This is not the case for reaching even where the average position was the same (Fig. 6E, upper panel). Although the correlation matrix was also markedly similar to that during latency (see Fig. 5A), the original body configuration was not recovered (Fig. 6E in the lower-left and right panels) because various reaching movements were included, which could re-validate the change point estimation.

Although the correlation matrix between 2- and 3-year-olds was congruent (Fig. 6BD, reprinted from 5BC), there was a statistical difference in predicting their touch errors (overlaid asterisks). The correlation matrix was vectorized and entered into a linear regression model to predict the touch error distance. For 2-year-olds, the latency matrix as the assembled dataset (regardless of the participants or target positions at this stage) can predict touch error (*F*(30, 436) = 1.666, *p* = .016). By contrast, for 3-year-olds, the reaching matrix can predict touch error (*F*(30, 308) = 1.636, *p* = .022). However, the latency matrix for 3-year-olds or the reaching matrix for 2-year-olds cannot predict their touch errors (*F*(30, 288) = 1.475, *p* = .057; *F*(30, 435) = 1.19, *p* = .229, respectively). For each model, the significant (and marginal for reference) variables are indicated by symbols (Fig. 6BD, *p* < .01**, *p* < .05*, *p* < .10.) with signs (positive or negative effects), which include the L-R upper body associations (i.e., hand movements) in the x coordinates regardless of age. Figure 6CF further depicts the association between body parts in tanglegram (R library “dendextend” (https://talgalili.github.io/dendextend/)) in terms of “correct” or “incorrect” labels using manual coding. Regarding latency (Fig. 6C), the incorrect matrix for 2-year-olds is characterized where upper (R)-lower (R) parts are associated (red ellipse), while the other tangle grams suggest that the upper (R) cluster was independently moving (yellow rectangle). Regarding reaching (Fig. 6F), the incorrect matrix for 3-year-olds is characterized where the upper (R)-lower (R) parts are associated again (red ellipse), while the other tangle grams suggest that the upper (R) and face clusters were associated in the x coordinates (yellow rectangle) (see Fig. S7A for the cross-correlation between the face and hand). If face movement is critical for precise touching (see Fig. 5D again), this could be related to their feedback/feedforward control in the AR mark test (Shen et al., 2010). This might have emerged as the weight between the latency/reaching duration as a function of feedback/feedforward control strategy between both age groups as follows.

### C. Internal Models and Their Weighting Changes

#### The Weight Between Feedback/Forward Strategy Changes Among Ages

Thus far, the Bodytoypo task mainly generated participants’ touch errors and the latency/reaching duration (Fig. 7B) as well as the association between body parts. These depend on the target location (see Fig. 5 and 6). This specifically varies the initial mark distance between the hand used and the target location when the mark appeared (Figure S6 for a descriptive correlation matrix among variables). Although categorical comparisons, such as 2- or 3-year-olds or correct/incorrect trials, have been effective in qualitatively depicting what was happening in the current task during the latency or reaching periods, the all-in-one statistics with a linear mixed model attempted to conclude these relationships continuously, using other covariates as mentioned above. Therefore, the best-fit model was explored using the R library (“lme4” (https://github.com/lme4/lme4/) and “lmerTest” (https://github.com/runehaubo/lmerTestR), where the initial inputs were latency duration, reaching duration, trial repetition, initial mark distance, and participants’ age in months as fixed effects, as well as participants’ ID and target labels as random effects; the touch error value was the output. This every-effect-model generated significant interaction effects (e.g., the interactions among reaching, age, repetition, and initial distance (*t*(422. 5) = 2.721, *p* = .0068)). The step function further identified the model with the minimum Akaike information criterion (AIC). This selected-effect-model still generated a significant interaction among latency/reaching, age, repetition, and initial distance (*t*(557. 0) = 2.30, *p* = .0218 for latency; *t*(554. 0) = 2.38, *p* = .0177 for reaching). Figure 7A presents the relationship between latency, reaching, and age in months in terms of predicting the touch errors (see Fig. S7B for other interactions when total RT was used instead of latency/reaching duration).

For younger months, both latency and reaching should be longer for correct touch (minimized touch error). However, generally, there was a robust anti-correlation between latency and reaching duration (Fig. S6). For older months, however, these should be shorter for correct touch. This might imply that younger months rely on slower feedback control when correct. Accordingly, if they administer a quick movement without visual feedback (i.e., ballistic exploration), they might make mistakes (see Fig. 7C; as shown below, the latency ratio did not vary for incorrect trials for 2-year-olds). Several months later, they no longer exhibited ballistic movements. The older participants may rely on feedforward control with a shorter latency/reaching when correct. Accordingly, if they exhibit longer reaching as feedback control, they may mistouch. This finding may conflict with a conventional understanding of feedback/feedforward weight with development [29]. Given that visually guided reaching through a mirror (e.g., mirror drawing [30]) can confuse even an adult; more weight on feedback control for the AR mark test is not the optimal strategy. Indeed, the trial repetition factor suggests that, for older months, the repetition increased their touch errors, presumably owing to habituation-driven exploration or playing with feedback control (Fig. S7B, the upper panels).

Figure 7B shows the weight between latency/reaching duration in relation to the touch error and correct/incorrect labels. During a reach, the duration distribution peaked for two-year-olds in incorrect trials and peaked for three-year-olds in correct trials. When the relative duration of latency (the ratio for total RT) was summarized between two- and three-year-olds (Fig. 7C, a contour plot for visualization), the target-dependent variance of the latency ratio might be observed for correct trials with 2-year-olds, and incorrect trials with 3-year-olds, which could be evaluating their feedback weight. However, for the correct trials among 3-year-olds, the initial mark distance was positively correlated with the latency ratio (Fig. S6) and interacted with the total RT for touch errors (Fig. S7B, the lower panels). The hand was synchronized with face movements in y-coordinates (Fig. S7A), which could be evaluated by their feedforward weight. The best strategy for the Bodytoypo task should be the optimal integration of feedback/feedforward control depending on the target location (e.g., initial mark distance). The current results suggest a developmental weight shift from ballistic explorative movements (incorrect trials) to feedback-based slow control (correct trials) for 2-year-olds and from feedback-based slow control (incorrect trials) to feedforward-based fast control (correct trials) for 3-year-olds. The current task could reveal the complex but optimal weight between the internal feedforward output, as well as the external feedback input. We further re-considered visual observations over the original videos, according to the statistical results.

#### Reaching Strategies and Evaluating Weights

Automatic analysis revealed developmental changes in toddlers’ reaching strategies: ballistic search, feedback control, and feedforward control. Ballistic search refers to the participant reaching for the mark quickly, without aiming (Supplementary Video S1). Feedback control is a slower reaching strategy in which hand movements are adjusted while watching the video monitor or actual body part (Supplementary Video S2). Lastly, feedforward control is a strategy for reaching the target quickly while anticipating localization (Supplementary Video S3).

To examine the validity of this categorization, we tested whether the three reaching strategies found in the automatic analysis were classifiable to human observers: two coders reviewed the video and classified all trials into one of the three strategies. An agreement was calculated using 32.5% of the total trials. However, the agreement was low (*κ* = .33), especially in the distinction between the ballistic exploration and feedforward, and between feedforward and feedback; a difference of approximately 25% in judgment. This result suggests that classifying reaching strategies by eye is inherently difficult. Children sometimes switch strategies within a single trial (Supplementary Video S4), making discrete categorization unreliable. Such ambiguity is better understood not as an exclusive choice among three strategies but as online reweighting of internal models, a process that our automated analysis can quantify by estimating the degree of contribution of each strategy. At the same time, a qualitative re-evaluation of the automated output revealed meaningful differences in hand posture during reaching. In particular, the use of the index finger emerged as a potential developmental indicator of changes in reaching strategies (Fig. S3).

## V. Discussion

The present study had two aims: (1) to evaluate whether the cross-reality mark task (Bodytoypo) serves as an effective measure of toddlers’ body-part localization and reaching trajectories, and (2) to examine whether these behavioral indices can capture developmental changes in visuo-proprioceptive integration. To address these aims, we analyzed not only correct and incorrect localization but also touch error (distance) embedded in the first touch, estimating the factors predicting touch error.

Regarding the validity of the task, most two- and three-year-olds were able to complete all trials, demonstrating that Bodytoypo is feasible and repeatedly applicable for toddlers. To explore the relation between this new task and body semantics, we evaluated correlations with parent-reported body-part vocabulary. No correlation was observed between touch-error rates and vocabulary scores for each body part. Consistent with Schwoebel and Coslett [8], this finding suggests that body semantics and body topology may constitute partially independent components of body representation. However, when body parts were categorized according to the presence or absence of lexical labels, unlabeled body parts consistently showed higher localization error across ages. This indicates that body semantics and body topology cannot be regarded as entirely independent. Furthermore, interaction patterns observed for joint and highly movable regions suggest that bodily understanding acquired through motor experience may contribute to localization performance.

The technical contributions of the task are twofold. First, the integration of behavioral experimentation with advanced sensing technology allowed real-time, markerless tracking of toddlers’ whole-body movements using OpenPose, which estimates posture from 2D images through deep learning and achieves detection quality comparable to adults [31], [32]. Given that studies using 3D markers report substantially higher attrition due to marker removal—for example, 31.8% in a previous study [33]—markerless tracking provides a substantial methodological advantage. Although AR marks were presented in near real time, touch judgments were made manually to avoid misclassification by 2D image processing, ensuring that toddlers correctly learned the rule that touching the virtual mark yields a reward.

Second, we automated trajectory analysis using machine-learning methods. Developmental research has historically relied on human classification of children’s behaviors, yet multiple strategies frequently coexist within a single action sequence, including the first touch. The present analysis does not aim to categorize discrete actions but instead quantifies how the weighting of reaching strategies changes developmentally. Thus, this work represents not a simple digitization of the mark test but a comprehensive integration of engineering (online task processing), machine-learning statistics (offline whole-body movement analysis), and developmental psychology (theoretical interpretation and experimentation).

Turning to the indicators of body representation derived from reaching trajectories, the developmental shift from ballistic exploration to feedback and feedforward control between two and three years offers important insights into the acquisition of body representation, particularly the role of visual information in sensorimotor control. Ballistic exploration relies more heavily on proprioception, whereas feedback control requires online visual adjustments based on an acquired body representation, and feedforward control emphasizes motor prediction [34]. Three-year-olds showed greater weighting toward feedforward control in correct reaches. Although localization while observing a mirror image can be confusing even for adults, the use of the index finger observed during feedforward control may reflect appropriate motor prediction, suggesting increasing integration among proprioception, visual information, and motor prediction. These weightings adapt flexibly to task demands, and the shorter latencies and reaching durations observed among three-year-olds indicate rapid adaptation facilitated by motor prediction.

Moreover, region-specific patterns observed in this study— such as the relative ease of reaching to facial and joint regions and the higher error rates for body parts without lexical labels— suggest that visuo-proprioceptive integration may mature at different rates across different body regions. This supports the introductory claim that developmental coordination between the sensorimotor body schema and visually based body topology is spatially heterogeneous rather than uniform.

As noted in the Introduction, prior research has emphasized the specific role of visual information in multisensory integration. Studies tracing the development of self-reaching show that infants younger than six months do not reach for a target while visually fixating it [17]. Research using the rubber hand illusion has likewise demonstrated that children rely more strongly on visual information than adults, responding preferentially to the visible hand [35]. These findings suggest that integrating visual information into proprioceptive processing requires both time and motor experience. Clarifying the developmental changes in this interaction remains an important task for future research.

This study also has limitations. Because Bodytoypo relies on real-time skeleton detection, the AR display involves approximately 0.33 seconds (around ten frames) of latency. Although this latency did not appear to confuse toddlers during task performance, it is undeniably large for applications requiring high-precision reaching. Continued technological improvements may reduce this latency, enabling application to more demanding motor tasks. Furthermore, because the present sample consisted only of Japanese toddlers, it remains difficult to generalize the observed independence between vocabulary acquisition and localization performance. Cross-cultural work is needed, and because Bodytoypo does not require verbal instruction, it is well-suited for studies across diverse cultural and linguistic contexts—even potentially across species.

Finally, tracking the development of body-part localization through mirror-based and self-image-based contexts may offer novel insights into early body-representation development. Although adults take mirror correspondence for granted, children do not initially treat the mirror image as self; yet by around 18 months, mirror self-recognition emerges naturally. The mirror serves as a simple yet powerful means of externalizing bodily self-information without relying on illusions such as the rubber hand or embodied avatars. It also constitutes an early extension of the minimal self by simultaneously supporting the emergence of body ownership and the sense of agency. Examining how localization accuracy across the whole body improves after the onset of mirror self-recognition may clarify the developmental trajectory of body topology in mirror contexts. Future extensions of Bodytoypo— such as implementing temporal delays or presenting distorted self-images with unnaturally extended limbs—may further illuminate how the malleability of body representation observed in adults arises in early development.

## Acknowledgment

We thank Ayano Ryoke and Moe Kobayashi for their help with the data collection and analysis. We also thank all caregivers and children who participated in this study. This manuscript has been deposited in preprint servers in bioRxiv. This work was supported by JSPS KAKENHI (Grant Numbers 19H04019, 23H03704) to Michiko Miyazaki, Tomohisa Asai, and Ryoko Mugitani.

## References

[1] F. J. Varela, E. Thompson, and E. Rosch, The Embodied Mind. MIT Press, 1991/2017.

[2] A. Noë, Action in Perception. MIT Press, 2004.

[3] S. Gallagher, How the Body Shapes the Mind. Oxford Univ. Press, 2005.

[4] M. Tsakiris, “The multisensory basis of the self: From body to identity to others,” Q. J. Exp. Psychol., vol. 70, no. 4, pp. 597–609, 2017.

[5] A. Maselli and M. Slater, “The building blocks of the full body ownership illusion,” Front. Hum. Neurosci., vol. 7, p. 83, 2013.

[6] F. de Vignemont, “Body schema and body image—Pros and cons,” Neuropsychologia, 2010.

[7] S. Gallagher and J. Cole, “Body image and body schema in a deafferented subject,” J. Mind Behav., pp. 369–389, 1995.

[8] J. Schwoebel and H. B. Coslett, “Evidence for multiple, distinct representations of the human body,” J. Cogn. Neurosci., vol. 17, no. 4, pp. 543–553, 2005.

[9] V. Slaughter et al., “Origins and early development of human body knowledge,” Monogr. Soc. Res. Child Dev., pp. 1–113, 2004.

[10] V. Camões-Costa, M. Erjavec, and P. J. Horne, “Comprehension and production of body part labels in 2-to 3-year-old children,” Br. J. Dev. Psychol., vol. 29, pp. 552–571, 2011.

[11] W. Waugh and C. Brownell, “Development of body-part vocabulary in toddlers,” Early Child Dev. Care, vol. 185, no. 7, pp. 1166–1179, 2015.

[12] A. Witt, S. Cermak, and W. Coster, “Body part identification in 1-to 2-year-old children,” Am. J. Occup. Ther., vol. 44, no. 2, pp.147–153, 1990.

[13] C. A. Brownell, S. Zerwas, and G. B. Ramani, “‘So big’: the development of body self-awareness in toddlers,” Child Dev., vol. 78, no. 5, pp. 1426–1440, 2007.

[14] C. A. Brownell et al., “When do toddlers represent their own body topography?,” Child Dev., vol. 81, pp. 797–810, 2010.

[15] A. J. Bremner et al., “Spatial localization of touch in the first year of life,” J. Exp. Psychol. Gen., vol. 137, pp. 149–162, 2008.

[16] C. C. de Klerk, M. L. Filippetti, and S. Rigato, “The development of body representations,” Proc. R. Soc. B, vol. 288, 2021.

[17] E. Somogyi et al., “Which limb is it? Responses to vibrotactile stimulation in early infancy,” Br. J. Dev. Psychol., vol. 36, no. 3, pp. 384–401, 2018.

[18] M. Botvinick and J. Cohen, “Rubber hands ‘feel’ touch that eyes see,” Nature, vol. 391, p. 756, 1998.

[19] D. Cowie, T. R. Makin, and A. J. Bremner, “Children’s responses to the rubber-hand illusion,” Psychol. Sci., vol. 24, pp. 762–769, 2013.

[20] D. Cowie, S. Sterling, and A. J. Bremner, “Development of multisensory body representation,” J. Exp. Child Psychol., vol. 142, pp. 230–238, 2016.

[21] Lee, L., Ma, W., & Kammers, M. (2021). The rubber hand illusion in children: What are we measuring?. Behavior Research Methods, 53(6), 2615–2630.

[22] J. A. Gottwald et al., “Body representation in childhood,” Dev. Sci., vol. 24, 2021.

[23] G. G. Gallup Jr., “Chimpanzees: self-recognition,” Science, vol. 167, pp. 86–87, 1970.

[24] B. Amsterdam, “Mirror self-image reactions before age two,” Dev. Psychobiol., vol. 5, pp. 297–305, 1972.

[25] M. Miyazaki and K. Hiraki, “Does rear-search error in the mark test indicate a uniqueness of body-representation in young children?” presented at the The sixth annual Budapest CEU Conference on Cognitive Development, January 7 2017. [Online]. Available: https://www.bcccd.org/down/bcccd_final_program_2017.pdf

[26] S. Imaizumi, T. Asai, and M. Miyazaki, “Cross-referenced body and action for the unified self: empirical, developmental, and clinical perspectives,” Body Schema and Body Image: New Directions, p. 194, 2021.

[27] M. Miyazaki, T. Asai, and R. Mugitani, “Touching!: An augmented reality system for unveiling face topography in very young children,” Front. Hum. Neurosci., vol. 13, 2019.

[28] Z. Cao, G. Hidalgo, T. Simon, S.-E. Wei, and Y. Sheikh, “OpenPose: Realtime multi-person 2D pose estimation using Part Affinity Fields,” IEEE Trans. Pattern Anal. Mach. Intell., vol. 43, no. 1, pp. 172–186, Jan. 2021, doi: 10.1109/TPAMI.2019.2929257.

[29] E. W. Bushnell, “The decline of visually guided reaching during infancy,” Infant Behav. Dev., vol. 8, no. 2, pp. 139–155, Apr. 1985, doi: 10.1016/S0163-6383(85)80002-3.

[30] D. Starch, “A demonstration of the trial and error method of learning,” Psychol. Bull., vol. 7, no. 1, p. 20, 1910, [Online]. Available: https://psycnet.apa.org/journals/bul/7/1/20/

[31] O. Ossmy and K. E. Adolph, “Real-Time Assembly of Coordination Patterns in Human Infants,” Curr. Biol., vol. 30, no. 23, pp. 4553–4562.e4, Dec. 2020, doi: 10.1016/j.cub.2020.08.073.

[32] S. Rocha and C. Addyman, “Assessing Sensorimotor Synchronisation in Toddlers Using the Lookit Online Experiment Platform and Automated Movement Extraction,” Front. Psychol., vol. 13, p. 897230, Jun. 2022, doi: 10.3389/fpsyg.2022.897230.

[33] B. A. Kahrs, W. P. Jung, and J. J. Lockman, “When does tool use become distinctively human? Hammering in young children,” Child Dev., vol. 85, no. 3, pp. 1050–1061, May 2014, doi: 10.1111/cdev.12179.

[34] U. Sailer, J. Randall Flanagan, and R. S. Johansson, “Eye–Hand Coordination during Learning of a Novel Visuomotor Task,” J. Neurosci., vol. 25, no. 39, pp. 8833–8842, Sep. 2005, doi: 10.1523/JNEUROSCI.2658-05.2005.

[35] D. Cowie, A. McKenna, A. J. Bremner, and J. E. Aspell, “The development of bodily self-consciousness: changing responses to the Full Body Illusion in childhood,” Developmental Science, vol. 21, no. 3. p. e12557, 2018. doi: 10.1111/desc.12557.

