## Supplemental_materials for "Tracking Action in Cross-Reality Mark Test Reveals Developing Body Representation Among Toddlers"

**This file includes:**

Figures S1 to S7

A

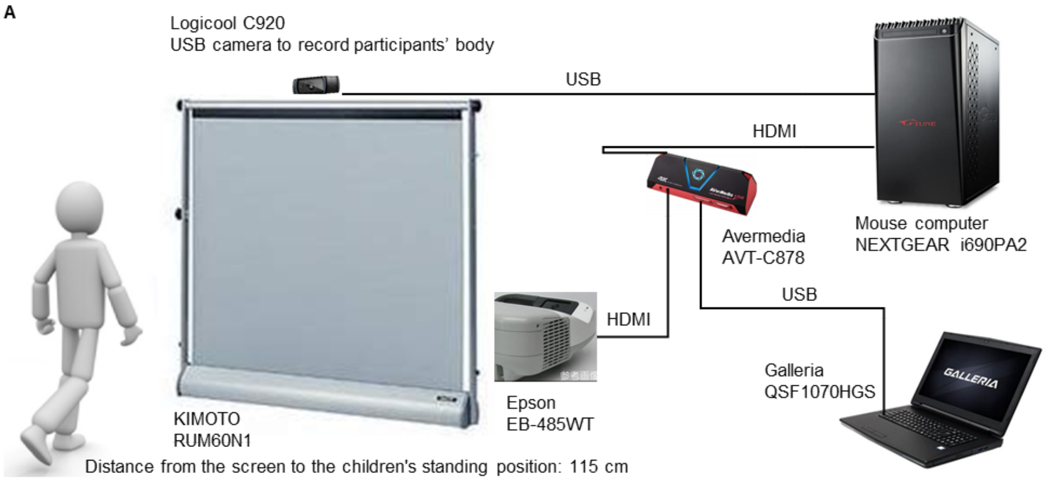


B

| Clusters | AI analysis(off-line) | | Bodytoypo(on-line) | |
|---|---|---|---|---|
| Face | | | upper forehead | (26) |
| | forehead | 1 | forehead | (27) |
| | eye(R/L) | 4, 2 | eye(R/L) | (15), (16) |
| | ear(R/L) | 5, 3 | ear(R/L) | (17), (18) |
| | nose | 6 | nose | (0) |
| | | | cheek(R/L) | (28), (29) |
| | | | mouse | (30) |
| | | | chin | (31) |
| | neck | 7 | | |
| upper body (R/L) | | | collarbone(R/L) | (41), (42) |
| | shoulder(R/L) | 14, 8 | shoulder(R/L) | (2), (5) |
| | | | upper arm(R/L) | (32), (33) |
| | elbow(R/L) | 15, 9 | elbow(R/L) | (3), (6) |
| | | | forearm(R/L) | (34), (35) |
| | wrist(R/L) | 16, 10 | | |
| | hand(R/L) | 17, 11 | hand(R/L) | (36), (37) |
| | thumb(R/L) | 18, 12 | | |
| | fingertip(R/L) | 19, 13 | | |
| Trunk | shoulder(M) | 20 | | |
| | | | navel | (40) |
| | hip(M) | 22 | | |
| lower body (R/L) | hip(R/L) | 27, 23 | | |
| | | | thigh(R/L) | (38), (39) |
| | knee(R/L) | 28, 24 | knee(R/L) | (10), (13) |
| | ankle(R/L) | 29, 25 | | |
| | toe(R/L) | 30, 26 | big toe(R/L) | (22), (19) |

C

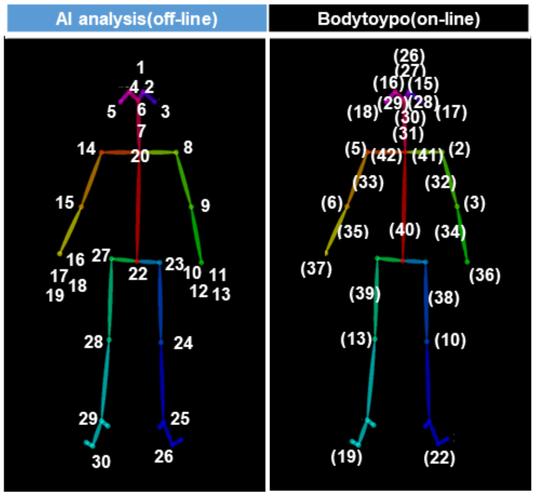


**Fig. S1. Apparatus for Bodytoypo and body parts used for presentation (online) and skeletal points used for analysis (offline).**

Note: (A) The apparatus for online task administration and its video recording for offline analysis. (B) Correspondence table of the list of 30 skeletal points used for analysis in the offline coding and the list of 30 body parts presented in Bodytoypo. (M) means midline. Body coordination was analyzed based on six body clusters, including right and left (R/L).

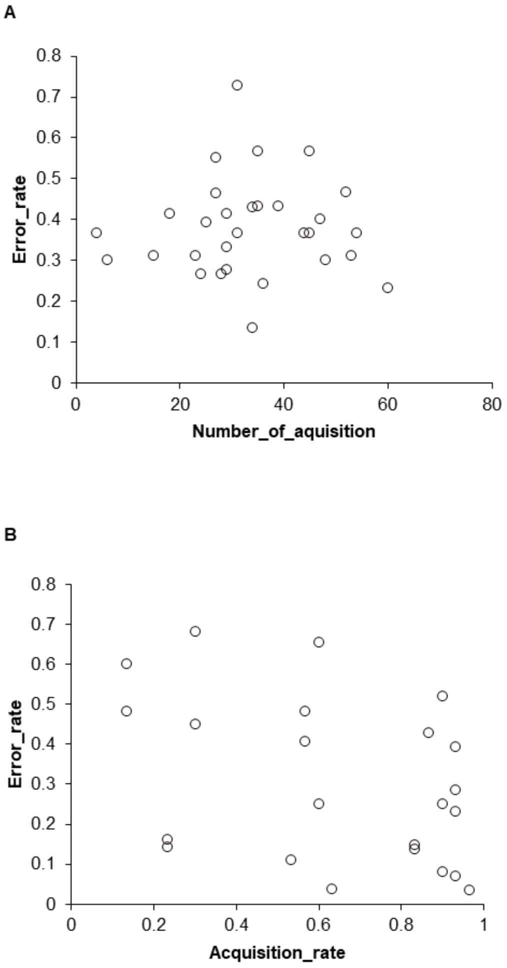


C

| | 2-yr | 3-yr | | 2-yr | 3-yr |
|---|---|---|---|---|---|
| Body | 50% | 83% | Tummy | 50% | 42% |
| Naked | 61% | 83% | Navel | 83% | 92% |
| Skin | 0% | 25% | Armpit | 33% | 75% |
| Head | 94% | 100% | Arm | 44% | 75% |
| Hair | 89% | 100% | Elbow | 11% | 58% |
| Down | 67% | 75% | Nail | 78% | 92% |
| Forehead | 56% | 75% | Hand | 89% | 92% |
| Eyebrow | 39% | 75% | Wrist | 6% | 42% |
| Eyelash | 17% | 67% | Finger | 89% | 92% |
| Eyelid | 0% | 33% | Thumb | 28% | 58% |
| Eye | 94% | 92% | Index finger | 22% | 25% |
| Ear | 94% | 92% | Middle finger | 11% | 25% |
| Nose | 94% | 100% | Third finger | 11% | 25% |
| Cheek | 78% | 92% | Pinkie | 11% | 33% |
| Mouth | 89% | 92% | Hips | 89% | 92% |
| Lip | 17% | 33% | Penis | 56% | 50% |
| Tongue (child word) | 50% | 67% | Foot | 94% | 92% |
| Tongue | 0% | 25% | Thigh | 6% | 25% |
| Beard | 44% | 75% | Knee | 50% | 67% |
| Tooth | 78% | 100% | Sole | 28% | 67% |
| Bad tooth | 39% | 83% | Ankle | 6% | 33% |
| Chin | 39% | 75% | Heel | 28% | 58% |
| Face | 89% | 92% | Toe | 11% | 42% |
| Throat | 17% | 58% | Mole | 22% | 67% |
| Neck | 61% | 75% | Bone | 11% | 50% |
| Shoulder | 50% | 75% | Blood | 67% | 92% |
| Back | 78% | 92% | Beak | 11% | 42% |
| Breast | 94% | 92% | Tail | 83% | 92% |
| Nipple | 6% | 33% | Wing | 28% | 75% |
| Stomach | 89% | 92% | Horn | 44% | 67% |

**Fig. S2. Correlation analysis of Bodytoypo's error rate and vocabulary acquisition related to body parts.**

Note: (A) Scatterplot of Bodytoypo's error rates and the number of word acquisitions. That correlation was .021. The partial correlation, controlling for age in months was .148, which is also not significant ($t(27) = .775$, $p = .445$, *n.s.*, 95% CI [-.231, .488]). (B) Scatterplot of Bodytoypo's error rates and acquisition rates of vocabulary had used in Bodytoypo. Of the 30 body parts used as targets in Bodytoypo, 23 with body part names were extracted and correlations were calculated; the correlation was -.346, not significant ($t(21) = -1.690$, $p = .106$, 95% CI [-.664, .077]). (C) Mean acquisition rates (comprehension + production) of body-related words (60 words) in each age group, based on caregiver reports.

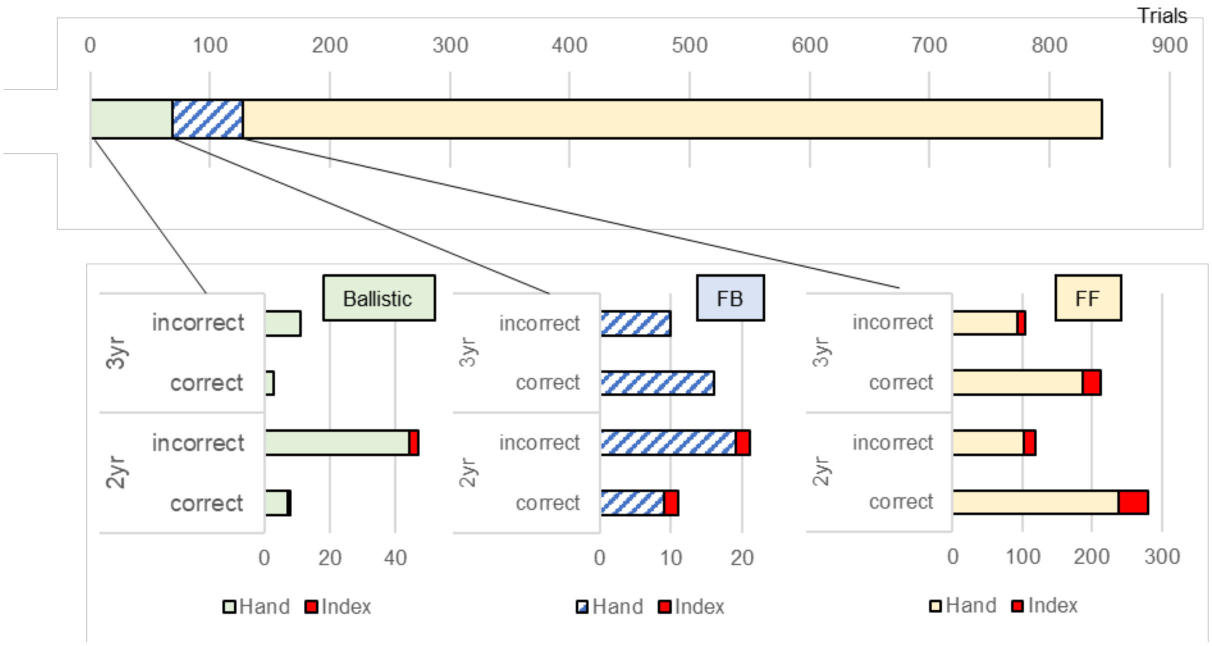


**Fig. S3. Developmental changes in using the index finger for each reaching strategy.**

Note: We focused on the shape of the hand during each reaching strategy and classified whether the hand (hand) or index finger (index) was used to reach or point. This is because we considered that hand shape includes differences that affect the accuracy of reaching localization. Specifically, since the hand (palm) touches a wider area, it is thought to increase the likelihood of correct touch even if the localization accuracy is not as high. On the other hand, pointing with the index finger is considered more difficult to localize. Classifying the hand shape, we found that 739 trials (87.56%) were reaching with the hand and 105 trials (12.44%) were pointing with the index finger. Focusing on index finger use trials, we found that 3-year-olds were shown to use the index finger only in the feedforward reaching; 2-year-olds used the index finger in all reaching strategies. This suggests that the 3-year-olds used the index finger only in reaching strategies that included prediction, i.e., reaching with some confidence.

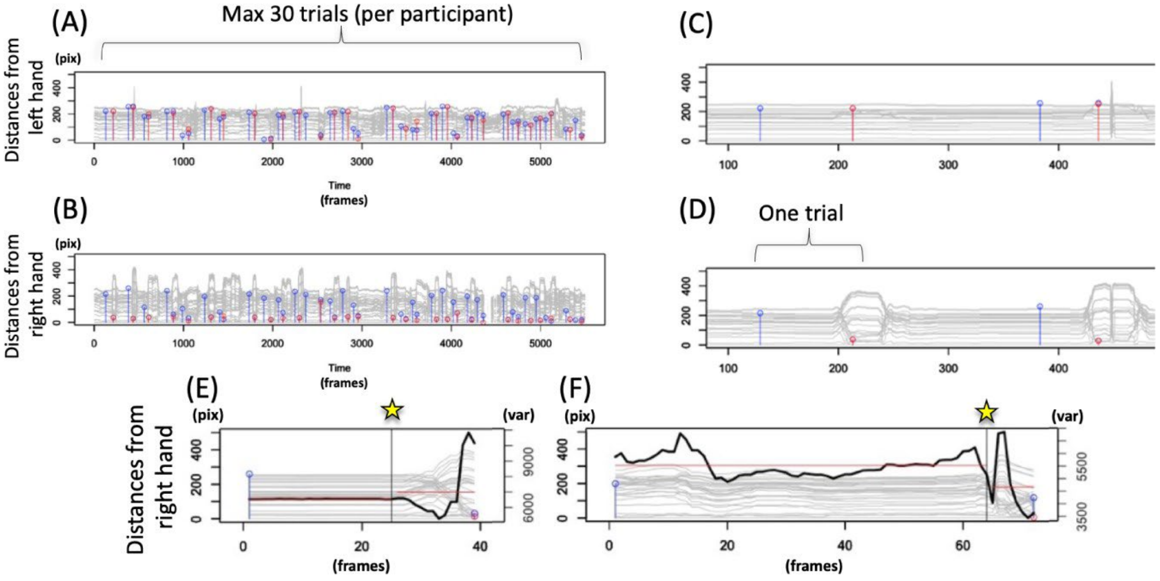


**Fig. S4. Typical trials from an exemplified child and the change point detection of the onset of reaching.**

Note: (A) Distance between the left hand and other parts (30 Gy lines). Blue circles indicate the mark timing and location (distance between the left hand and the mark). Red circles indicate the first-touch timing and touched location (distance between the left hand and the touched location). (B) Similarly, the distance between the right hand and other parts. (C, D) Partial expansions of A and B. The task lasted approximately 3-5 minutes with a maximum of 30 trials. In this example, the child used his or her right hand to touch the mark. Therefore, almost 30 “events” are observed in B, regardless of the location of the mark, while A is generally silent. In addition, the red circles were almost zero distances in B, which suggests right-hand touches because the touched location should be almost the same as the location of the hand used. In other words, non-zero red circles for the used hand indicate “incorrect” touches. (E, F) The vertical line with a star between the mark (blue circle) and the touch (red circle) timing is the estimated onset of reaching by a change-point detection algorithm for the time series of variance from 30 distances (black broken line). In an excellent example (E), where the child was almost “silent” till moves. Even in a delicate example (F), where the child was “vigorous,” the estimation seems to work in terms of visual inspection of distance changes (gray broken lines). This is because both the temporal mean (horizontal straight line) and temporal variance of the black broken line should be dramatically changed at the reaching onset. The globally harmonized movement until reaching, while the locally (the used hand) rapid movement during reaching should be observed even for vigorous children. The former can be called “response latency” while the latter is called “reaching duration”

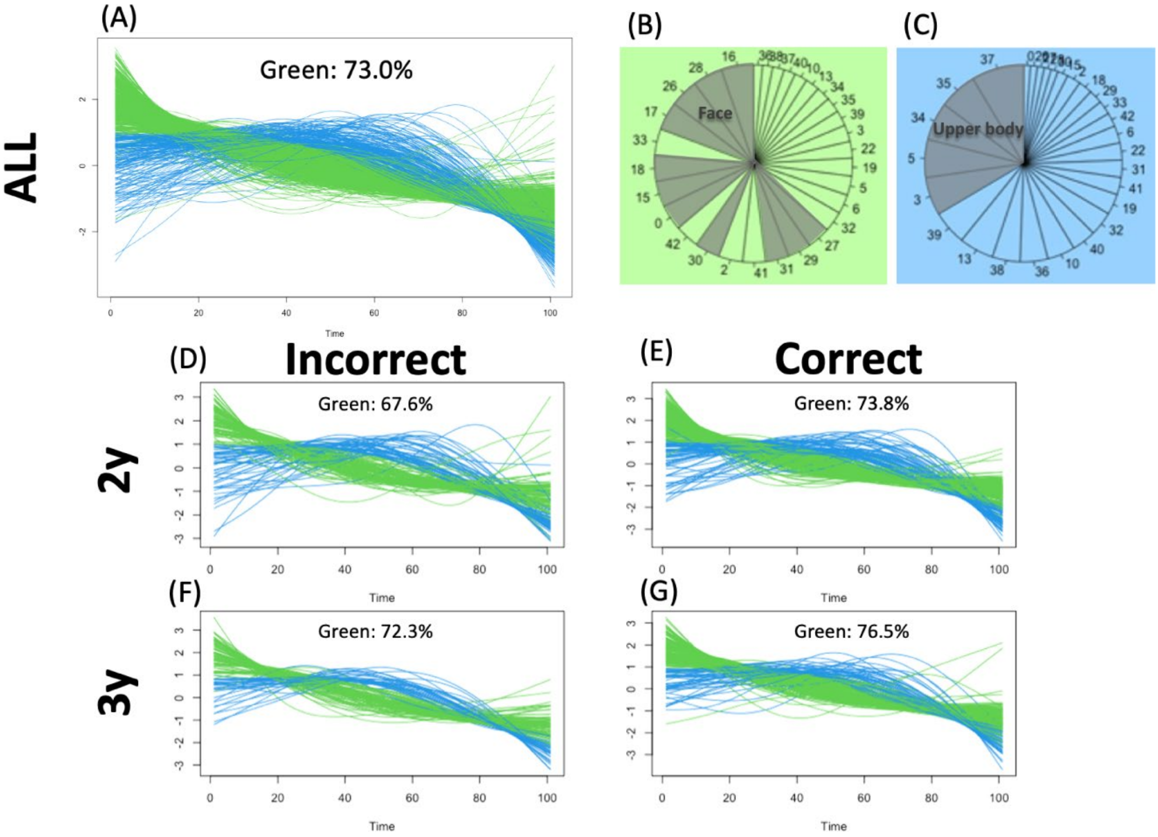


**Fig. S5. Proximate reaching trajectory in 2D in relation to the targeted body parts.**

Note: (A) The horizontal axis represents the normalized time elapsed for each trial, while the vertical axis represents the normalized hand-mark distance (the distance between the mark and the hand used). Each line represents each trial (i.e., a proxy for the trajectory). The initial hand-mark distance (at time 0 for the onset of reaching) was gradually reduced (at time 100 for touching) in general after temporal smoothing. However, depending on the target location, the 2D-projected trajectory from the camera can be categorized into clusters. The time-series clustering (the R library "TSclust") is divided into two trajectories: convex downward (green) or upward (blue). (B) The former trajectories mainly consist of the face-part target (shaded gray). (C) The latter consists of the upper-body-part target (shaded gray). Even when we compare among ages and in/correct trials, the congruent results emerge, which indicates that the reaching behavior itself should be basically the same among participants and trials (except for a bit more correct trials for face parts for 3-year-olds, for example).

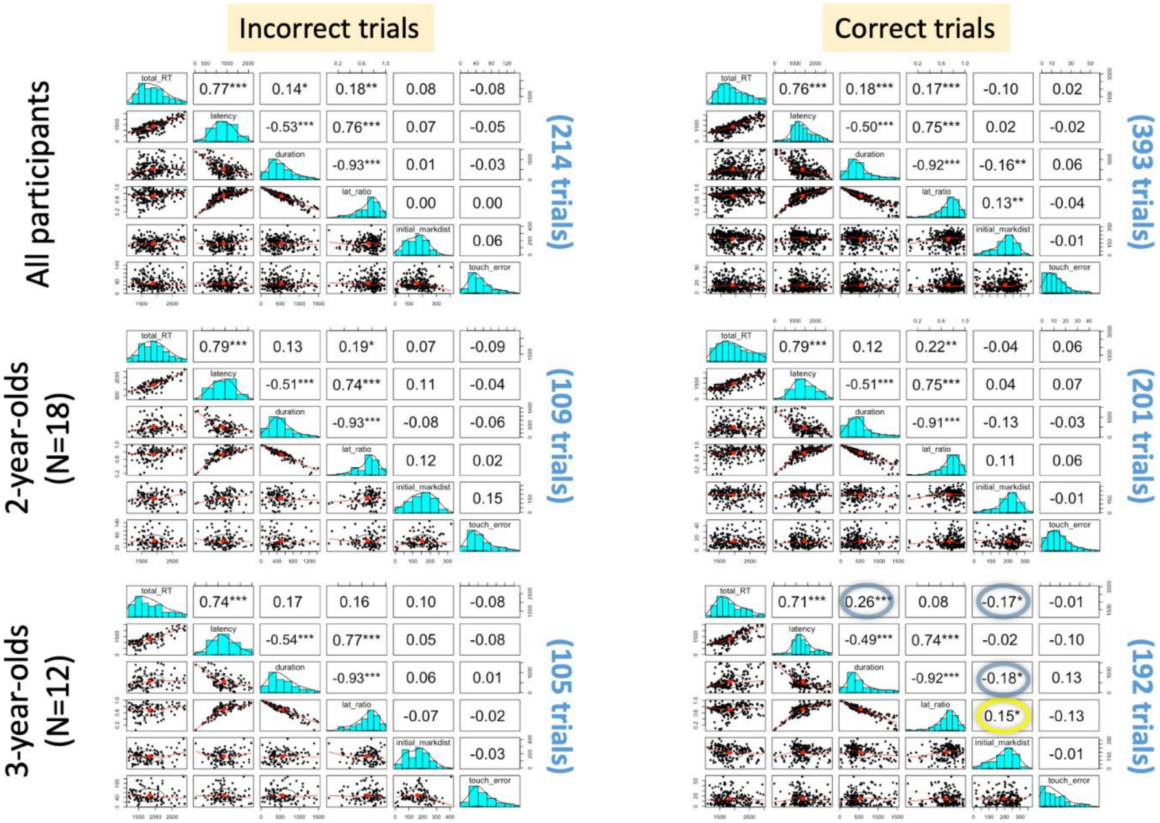


**Fig. S6. Descriptive correlation matrix among the estimated indices.**

Note: A total of 852 trials were completed by 30 children (ideal maximum was 30 children × 30 trials = 900 trials). After omitting unreliable trials (total RT < 1s, > 3s, or touch error > 150 pix), 607 trials were further analyzed. The omitting criteria were decided by a visual consultant to each distribution (on the diagonal, see also Figure 7AB). The main reason to be omitted was too long total RT (over 3s is approximately 3SD + mean for the omitted distribution) when children were not noticed by the mark, less motivated to touch, play, and temporally unattended, etc. The total RT was from the mark appearing to the first touch (frame-by-frame manual video coding). The response latency was until the estimated onset of reaching (using the machine-learning algorithm). The reaching duration was determined from the onset of the first touch. The latency ratio was the fraction of response latency to total RT. The initial mark distance was the Euclidean 2D distance between the mark and the hand used at the mark appearing (by frame-by-frame manual video coding). The touch error was the similar distance between the touched position and the mark at that moment (through frame-by-frame manual video coding). Manual video coding was conducted by multiple coders to avoid a specific coding policy (where the two coders were internals to decide the coding instruction and the two were requested naïve externals). Accordingly, manually trimmed and coded trial-wise video files (852 files) were further processed and analyzed into x-y time-series data for 30 body parts. See the main text for the following analysis:

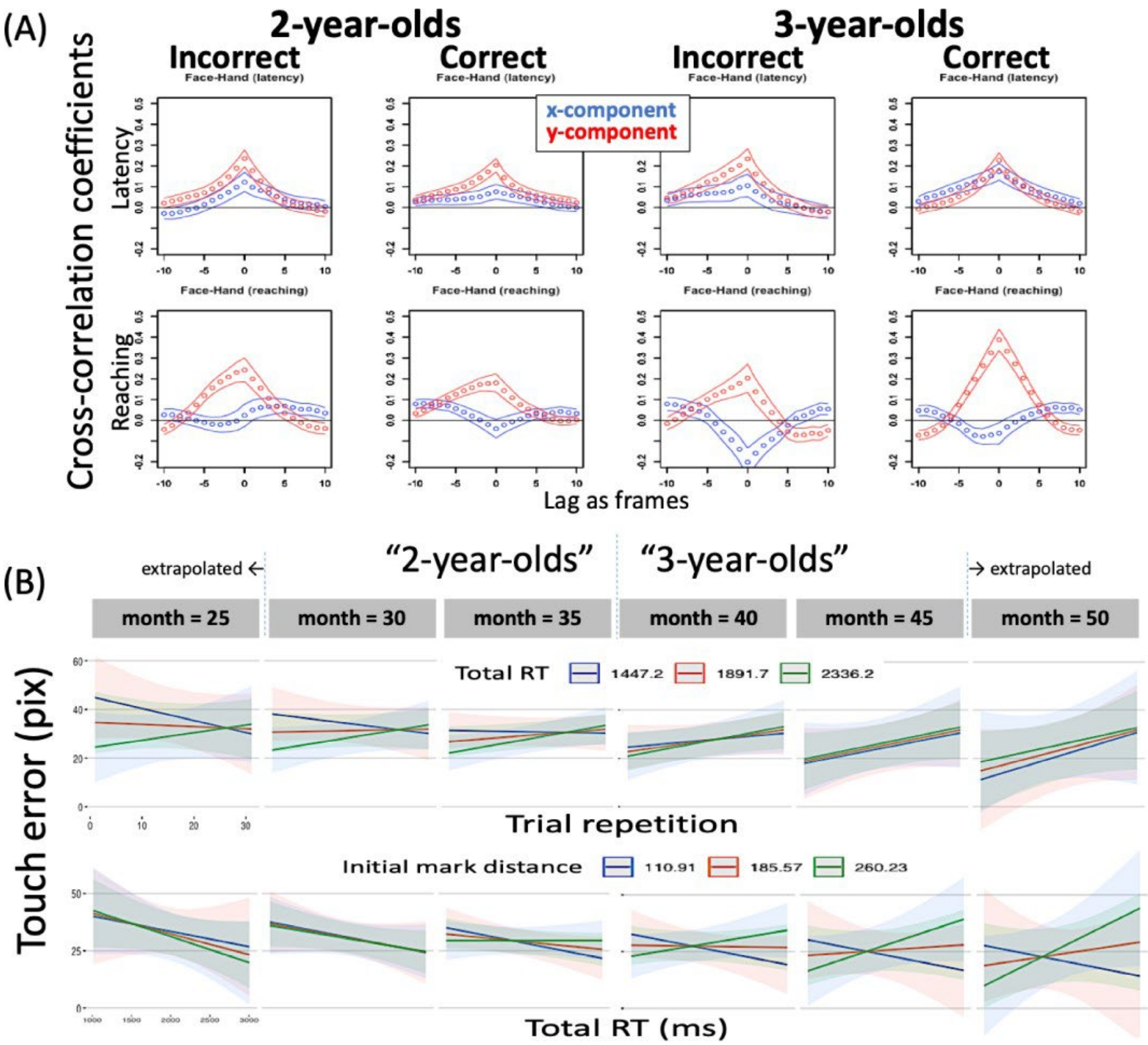


**Fig. S7. Cross-correlation between the face and hand (A) and the other LMM interactions (B).**

Note: During reaching, the face (or head) movement seems important for correct touches, especially for 3-year-olds. (A) The cross-correlation coefficients are plotted as a function of lag as frames (30 fps). If the peak is shifted toward the left (or right), the hand moves first (or the face moves first). For the current task, however, the peaks were almost on lag 0, suggesting that the face and hand were moved synergistically. This was robust for correct trials in y-coordinates and for incorrect trials in x-coordinates for 3-year-olds. (B) The relationship among total RT, trial repetition, initial distance, and age (months) for predicting touch errors is illustrated. In terms of trial repetition (upper panels), the older months exhibited larger touch errors with repetition. On the other hand, the initial mark distance varied their total RT for older months, while it did not affect younger months (the lower panels). See the main text for the potential interpretations of these.
